# Regulation of KIF1A motility via polyglutamylation of tubulin C-terminal tails

**DOI:** 10.1101/410860

**Authors:** Dominique V. Lessard, Oraya J. Zinder, Takashi Hotta, Kristen J. Verhey, Ryoma Ohi, Christopher L. Berger

**Author notes:** **Corresponding Author:** Christopher L. Berger, Ph.D., Department of Molecular Physiology & Biophysics, University of Vermont, 122 HSRF, 149 Beaumont Avenue, Burlington, Vermont 05405.

## Abstract

Axonal transport is a highly regulated cellular process responsible for site-specific neuronal cargo delivery. This process is mediated in part by KIF1A, a member of the kinesin-3 family of molecular motors. It is imperative that KIF1A’s highly efficient, superprocessive motility along microtubules is tightly regulated as misregulation of KIF1A cargo delivery is observed in many neurodegenerative diseases. However, the regulatory mechanisms responsible for KIF1A’s motility, and subsequent proper spatiotemporal cargo delivery, are largely unknown. One potential regulatory mechanism of KIF1A motility is through the posttranslational modifications (PTMs) of axonal microtubules. These PTMs, often occurring on the C-terminal tails of the microtubule tracks, act as molecular “traffic signals” helping to direct kinesin motor cargo delivery. Occurring on neuronal microtubules, C-terminal tail polygutamylation is known to be important for KIF1A cargo transport. KIF1A’s initial interaction with microtubule C-terminal tails is facilitated by the K-loop, a positively charged surface loop of the KIF1A motor domain. However, the K-loop’s role in KIF1A motility and response to perturbations in C-terminal tail polyglutamylation is underexplored. Using single-molecule imaging, we present evidence of KIF1A’s previously unreported pausing behavior on multiple microtubule structures. Further analysis revealed that these pauses link multiple processive segments together, contributing to KIF1A’s characteristic superprocessive run length. We further demonstrate that KIF1A pausing is mediated by a K-loop/polyglutamylated C-terminal tail interaction and is a regulatory mechanism of KIF1A motility. In summary, we introduce a new mechanism of KIF1A motility regulation, providing further insight into KIF1A’s role in axonal transport.

## INTRODUCTION

Axonal transport is a critical process for neuronal viability and function involving the highly choreographed long distance trafficking of cargo along the axon. This process is facilitated in part by members of the kinesin superfamily of motors that utilize the mechanochemical coupling of ATP-hydrolysis [1] to carry motor-specific cargo in the anterograde direction along axonal microtubules.

The neuron-specific kinesin-3 family member KIF1A [2] is a key mediator of axonal transport by spatiotemporally delivering neuronal cargo, such as dense core vesicles [3] and synaptic vesicle proteins [2], along axonal microtubules [4, 5]. Upon cargo-mediated dimerization and release of steric autoinhibition, KIF1A motors are reported to exhibit “superprocessive” motility behavior, traveling comparatively long distances on microtubules when equated to conventional kinesin motility [6, 7]. The discovery of this superprocessive behavior supports KIF1A as an important player in long distance axonal transport, providing further insight into KIF1A’s role in many neuronal processes such as synaptogenesis [8] and neurogenesis [9]. The importance of tightly regulated KIF1A cargo delivery is highlighted in neurodegenerative diseases where irregular KIF1A cargo accumulation is observed [10-12]. However, our understanding of this disease-state presentation is limited by the gap in knowledge of the mechanisms governing regulation of KIF1A motility.

Beyond serving as tracks for kinesin motors, microtubules help control motor specificity on select microtubule populations by delivering important directional cues. These “traffic signs” are frequently introduced via microtubule posttranslational modifications (PTMs), often on the C-terminal tails of microtubule surface [13, 14]. While there are many recent discoveries of how microtubule PTMs regulate different kinesin families [15-17], how this concept can be extended to kinesin-3 motors such as KIF1A is relatively unexplored. One potential PTM regulating KIF1A trafficking is the heavy polyglumylation of both α- and β-tubulin subunits in neurons [18] via the tubulin tyrosine ligase-like family of enzymes [19, 20]. KIF1A’s dependence on optimal levels of polyglutamylation has been observed in the ROSA22 mouse model, characterized by a loss of neuronal α-tubulin polyglutamylation leading to altered KIF1A cargo trafficking and reduced KIF1A affinity for microtubules [21]. These physiological repercussions of altered KIF1A function highlight the importance of further studying interactions between KIF1A and the polyglutamylation of C-terminal tails of tubulin at the molecular level.

Regulated levels of polyglutamylation may impact KIF1A motility via the lysine rich surface loop 12 in the KIF1A motor domain, known as the K-loop and characteristic of the kinesin-3 motor family [22, 23]. Previous work has demonstrated that the K-loop is a critical component for optimal KIF1A function. From a catalytic perspective, it is thought that the K-loop is necessary for KIF1A’s structural interaction with the microtubule during the ATP-hydrolysis cycle [24]. Furthermore, it is known that the K-loop electrostatically tethers to the glutamic-acid rich C-terminal tail of tubulin [25, 26], mediating processes such as KIF1A affinity for the microtubule surface [22]. This K-loop/C-terminal tail relationship, combined with the physiological detriment of reduced microtubule polyglutamylation, illustrates that specific interactions between the KIF1A K-loop and C-terminal tails of tubulin are essential for optimal KIF1A function. Therefore, we hypothesized that C-terminal tail polyglutamylation regulates KIF1A motility and behavior on microtubules mediated by interactions with the KIF1A K-loop structure. Using single-molecule total internal reflection fluorescence microscopy, we first tested our hypothesis by quantifying the motility and behavioral response of KIF1A on various microtubule lattices. We next went on to perturb the K-loop/C-terminal tail interaction. Our findings lead us to present the previously unreported pausing behavior of KIF1A on multiple microtubule lattices. Furthermore, we present that KIF1A’s pausing behavior and motility is reliant upon interactions with the microtubule C-terminal tails and regulated by tubulin polyglutamylation.

## RESULTS

### KIF1A exhibits pausing behavior and superprocessive motility on taxol-stabilized and GMPCPP microtubules assembled from brain tubulin

Initial single molecule TIRF microscopy observations revealed that KIF1A possesses the novel ability to undergo extensive pausing during a run on taxol-stabilized microtubules (Figure 1A, Movie S1). While other kinesin motors have been shown to exhibit transient pauses [27, 28], the high pause frequency (Figure 2E, Table 1) of KIF1A has not been previously characterized, compelling us to investigate this behavior further. Additionally, the discovery of this novel pausing behavior led us to redefine our nomenclature of KIF1A motility by dissecting KIF1A motility into three segments (Figure 1A). First, we define the *processive run length* as segments where, within a single event, KIF1A is moving at a constant velocity. Next, processive run lengths are connected by *pauses*, occurring at the beginning of processive runs, in between processive runs, or after processive runs. While representative KIF1A kymographs represent long pauses (Figure 1C, 2A, 2B), it is important to note that the majority of pausing events are less than 1.5 seconds and occur at stochastic locations on the microtubule (Figure 1C, Table 1). Lastly, we define the *overall run length* as the summation of processive run lengths and pauses during a single motility event on the microtubule. Of note, KIF1A pausing behavior (Figure 1B, 1C) is best revealed at low motor, single-molecule concentrations of KIF1A, as it can be obscured at high motor density on the microtubule surface.

**Table 1.** Summary of KIF1A motility and behavior on microtubules across all conditions. * = p<0.001 relative to KIF1A on taxol-stabilized MTs.

| | Overall Run Length ($\mu\text{m}$) | Overall Speed ($\mu\text{m/s}$) | Processive Run Length ($\mu\text{m}$) | Processive Speed ($\mu\text{m/s}$) | Pause Frequency (# pauses/overall run) | Pause Duration (sec) | Landing Rate (events/$\mu\text{m}/\text{sec}$) |
| --- | --- | --- | --- | --- | --- | --- | --- |
| **Taxol** | $6.24 \pm 2.09$ | $1.35 \pm 0.41$ | $2.95 \pm 1.07$ | $2.01 \pm 0.53$ | 0.95 | $2.25 \pm 2.99$ | $7.93 \pm 1.25$ |
| **GMPCPP** | $6.01 \pm 2.17$ | $1.27 \pm 0.28$ | $2.81 \pm 0.91$ | $2.06 \pm 0.54$ | 1.02 | $1.40 \pm 1.46$ | $4.86 \pm 0.88^*$ |
| **Subtilisin** | $3.31 \pm 1.34^*$ | $1.58 \pm 0.38$ | $2.91 \pm 1.16$ | $1.94 \pm 0.50$ | 0.18* | $1.23 \pm 0.68$ | $2.80 \pm 0.42^*$ |
| **Tri-Ala** | $2.41 \pm 1.42^*$ | $1.66 \pm 0.45$ | $2.00 \pm 1.11^*$ | $1.72 \pm 0.45$ | 0.23* | $1.01 \pm 0.53$ | $0.64 \pm 0.14^*$ |
| **HeLa Tubulin** | $4.12 \pm 1.48^*$ | $1.70 \pm 0.41$ | $2.98 \pm 0.96$ | $1.87 \pm 0.40$ | 0.40* | $1.23 \pm 1.46$ | $0.49 \pm 0.13^*$ |

\* = $p < 0.001$ relative to KIF1A on taxol-stabilized MTs.

**Figure 1.**
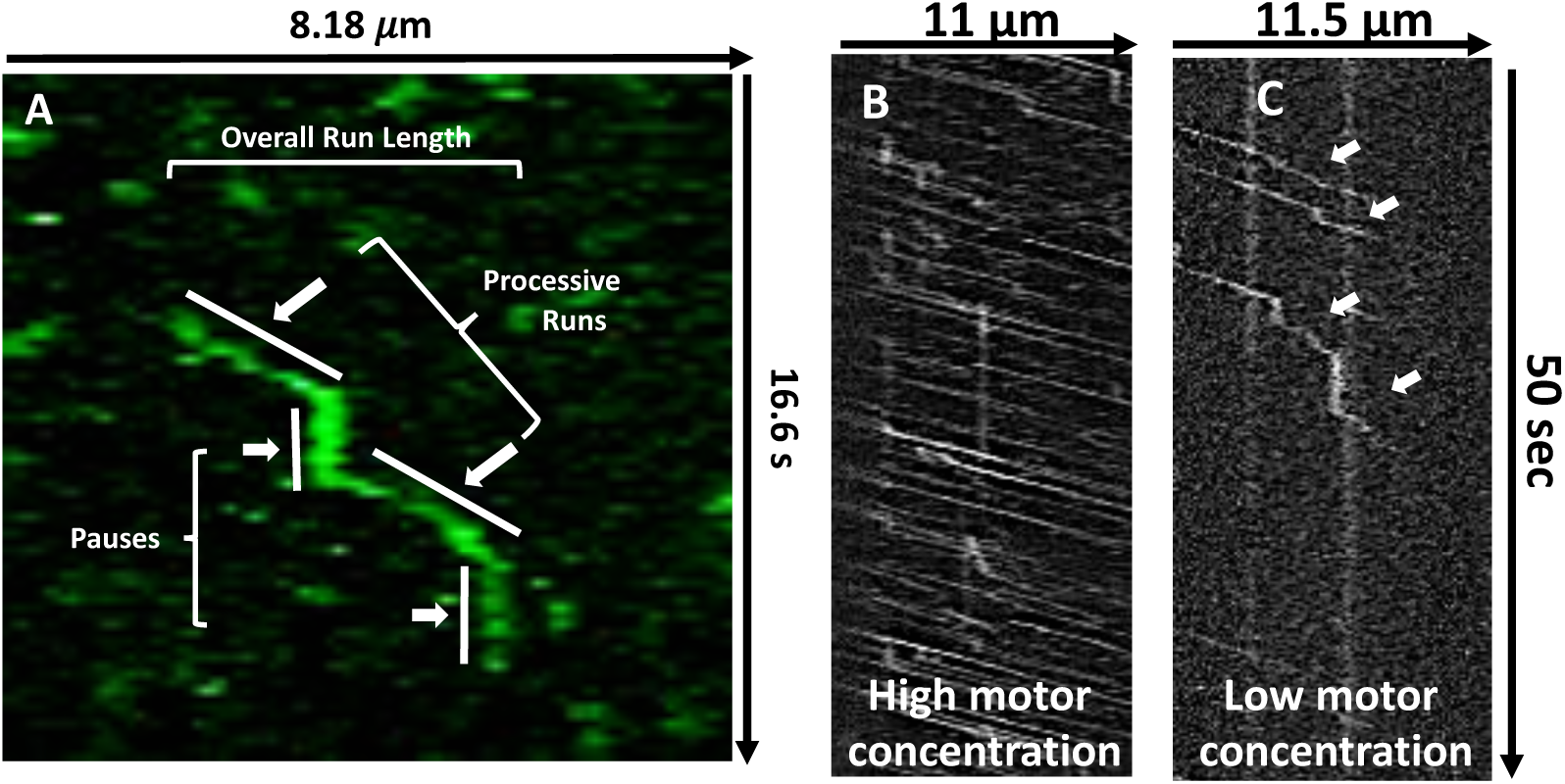
**A)** Representative kymograph with corresponding nomenclature used to describe novel KIF1A motility behavior. **B)** Representative kymographs detailing that at high motor concentrations pausing becomes obscured by surrounding motor motility while at **C)** low motor concentrations, extensive pausing behavior is revealed at stochastic positions on the microtubule (represented by white arrows).

**Figure 2.**
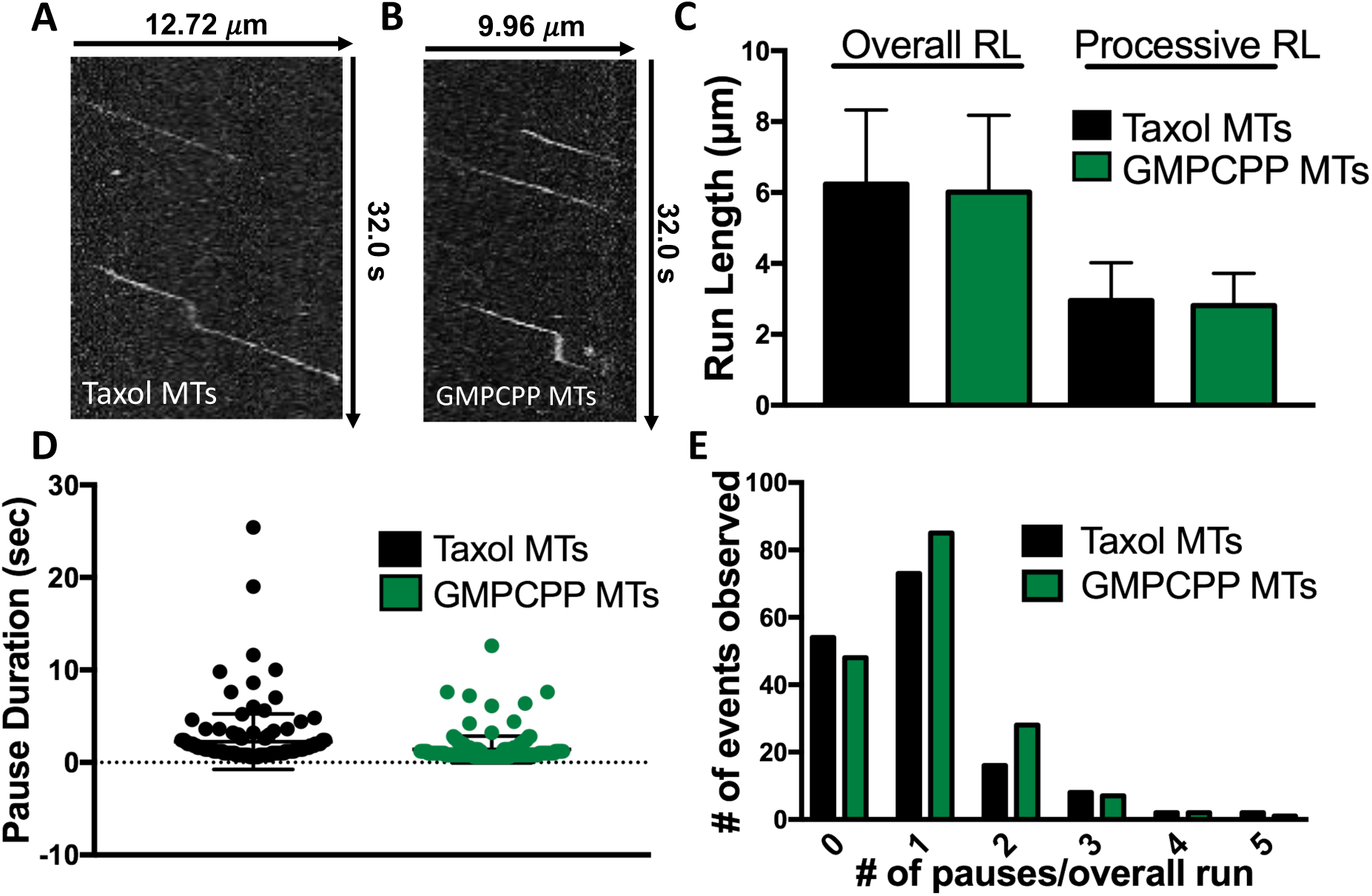
**A)** Representative kymograph detailing that KIF1A pausing behavior is maintained on taxol-stabilized and **B)** GMPCPP microtubules. **C)** On taxol-stabilized and GMPCPP microtubules, KIF1A overall run length (RL) (6.24 ± 2.09 µm [n=165] and 6.01 ± 2.17 µm [n=171], respectively) and processive run length (2.95 ± 1.07 µm [n=282], and 2.81 ± 0.61µm [n=305], respectively) are not significantly different. **D)** Duration of KIF1A pauses is not significantly different on taxol-stabilized (2.25 ± 2.99 sec [n=147]) and GMPCPP (1.40 ± 1.46 sec [n=152]) microtubules. **E)** KIF1A exhibits a similar pause frequency on taxol-stabilized (0.95 pauses/overall run [n=165]) and GMPCPP (1.02 pauses/overall run [n=171]) microtubules. Run length values and standard deviations were calculated as previously reported [73]. Kinesin-1 was used as an experimental control across all conditions (Figure S2-S5). Each condition is representative of at least four independent experiments. Mean ± SD reported.

To confirm that this behavior was not dependent on the facilitated dimerization of our leucine zipper construct (KIF1A-LZ-3xmCitrine), we repeated our single molecule experiments with a non-leucine zipper construct (KIF1A-GFP). We confirmed that in the absence of the leucine zipper, KIF1A exhibits extensive pausing behavior and superprocessive motility (Figure S1). Additionally, we observed an increase in purely diffusive events with this construct (Figure S1A), likely resulting from KIF1A monomers [7]. Our results demonstrate that the KIF1A-LZ-3xmCitrine construct is an appropriate model for dimeric KIF1A behavior. As we currently only want to investigate the processive, dimeric KIF1A motor, we performed all other conditions with the LZ-construct.

We next investigated KIF1A’s pausing by quantifying the above mentioned pausing and motility characteristics on both taxol-stabilized and GMPCPP microtubules, which are known to have more protofilaments [29, 30], and fewer lattice defects [31] than taxol-stabilized microtubules. KIF1A pausing was maintained on both microtubule lattices (Figure 2A, B), with pause durations of 2.25 ± 2.99 sec (Taxol) and 1.40 ± 1.46 sec (GMPCPP) and similar pause frequencies of 0.95 (Taxol) and 1.02 (GMPCPP) (Figure 2D, 2E, Table 1). In comparing taxol-stabilized and GMPCPP microtubules, we observed no significant decrease in overall run length (6.24 ± 2.09 µm and 6.01 ± 2.17 µm) or processive run length (2.95 ± 1.07 µm, and 2.81 ± 0.61µm), respectively (Figure 2A, Table 1). Additionally, we observed that the speed in which KIF1A processes along either taxol-stabilized or GMPCPP microtubules was consistent, with overall speed (1.35 ± 0.41 μm/s and 1.27 ± 0.28 μm/s, respectively; Table 1, Figure S2) and processive speed (2.01 ± 0.53 μm/s and 2.06 ± 0.54 μm/s, respectively; Table 1, Figure S3) being almost identical. While many of the pausing and motility characteristics of KIF1A were consistent across these two microtubule lattices, there was a significant decrease in KIF1A’s ability to land on GMPCPP microtubules in the ADP state, 4.86 ± 0.88 events/μm/min compared to 7.93 ± 1.25 events/μm/min on taxol-microtubules (Figure 6, Table 1). These results confirm that KIF1A’s unique pausing behavior is maintained across microtubules of varying protofilament and nucleotide composition.

### Interaction with tubulin C-terminal tails mediates KIF1A pausing behavior, landing rate and superprocessive motility

To investigate a potential mechanism facilitating KIF1A pausing, we considered the known interaction of KIF1A’s lysine-rich K-loop with tubulin’s glutamate-rich E-hook projections [22, 23]. Previous work has shown that the positively charged surface loop 12 (K-loop) of the KIF1A motor domain interacts with the microtubule C-terminal tails, helping to anchor KIF1A to the microtubule [25] and mediate KIF1A landing on the microtubule [22]. Additional cellular work has shown that loop 12 is important for regulation in KIF1A dendritic cargo sorting [32]. With this previously established interaction, we asked if subtilisin-mediated proteolytic removal [33] (Figure 3A) of taxol-stabilized microtubule C-terminal tails could result in regulation of KIF1A motility and pausing behavior. On subtilisin-treated microtubules, we observed a 47% reduction of KIF1A overall run length when compared to KIF1A on untreated taxol-stabilized microtubules (Figure 3B, 3C, Table 1). Subtilisin treatment of microtubules did not significantly change the processive run length of KIF1A (Figure 3C, Table 1), however we did observe a 37% decrease in pause duration and a 79% reduction in pause frequency (Figure 3D, 3E, Table 1). Resulting from fewer pausing events, KIF1A’s overall speed increased on subtilisin-treated microtubules from 1.35 ± 0.41 μm/s (untreated, taxol-stabilized microtubules) to 1.58 ± 0.38 μm/s (Table 1, Figure S2). However, processive speed remained relatively unchanged between untreated, taxol-stabilized, and subtilisin-treated microtubules (Table 1, Figure S3).

**Figure 3.**
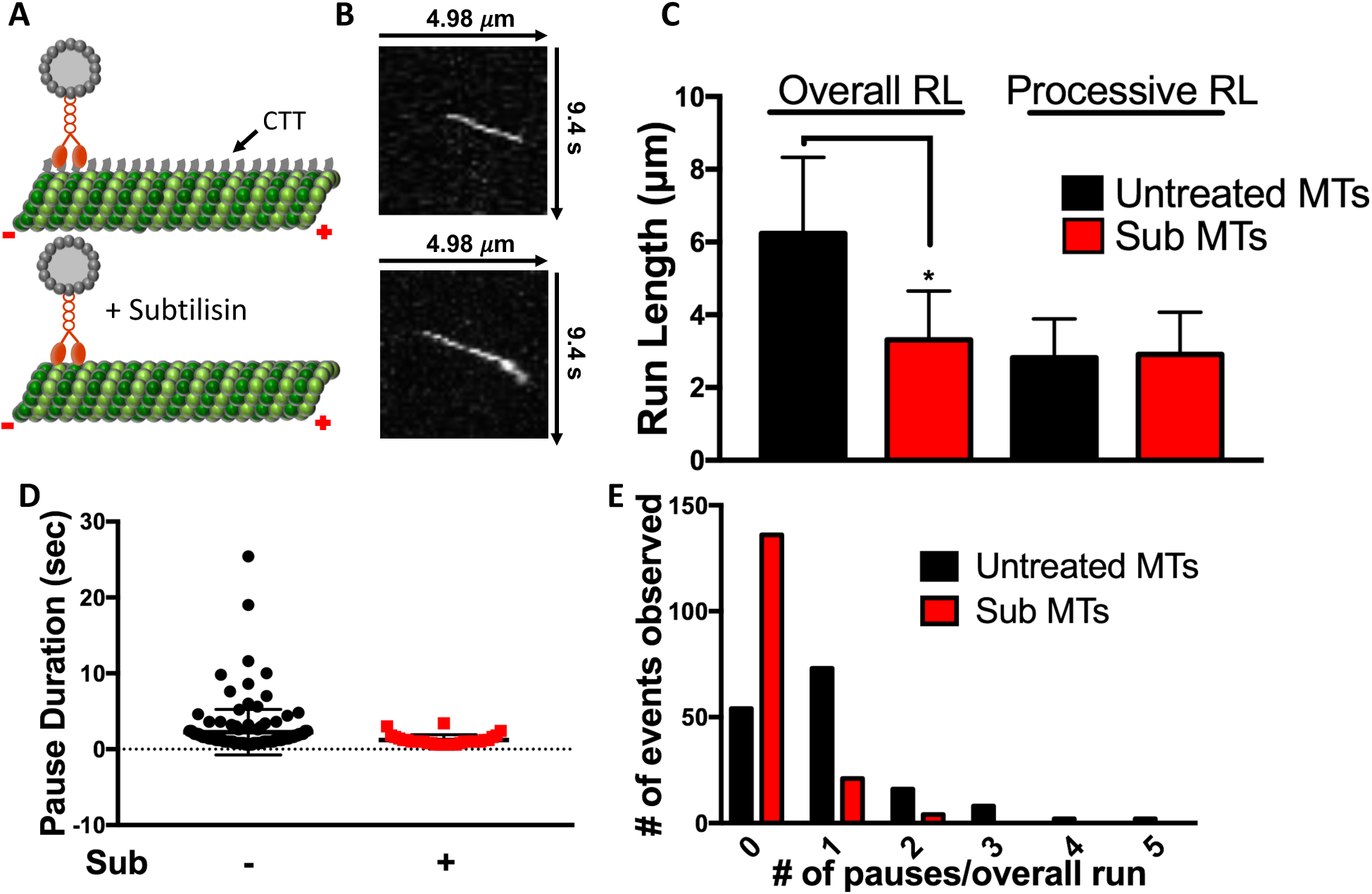
**A)** Cartoon depiction of paclitaxel-stabilized microtubule C-terminal tail (CTT) cleavage upon addition of subtilisin. **B)** Representative kymographs reveals that subtilisin (Sub) treatment, and subsequent removal of C-terminal tails, dramatically reduced KIF1A pause frequency and overall run length. **C)** The overall run length of KIF1A was reduced from 6.24 ± 2.09 µm to 3.31 ± 1.34 µm [n=171] upon subtilisin treatment of microtubules, while processive run length was not significantly changed (2.95 ± 1.07 µm and 2.91 ± 1.16 µm [n= 205], respectively). **D)** Pause duration decreased from 2.25 ± 2.99 sec to 1.23 ± 0.68 sec [n=34] on subtilisin-treated microtubules. **E)** KIF1A pause frequency was significantly decreased from 0.95 pauses/overall run to 0.18 pauses/overall run upon subtilisin-treatment of taxol-stabilized microtubules. Run length values and standard deviations were calculated as previously reported [73]. Kinesin-1 was used as an experimental control across all conditions (Figure S2-S5). Each condition is representative of at least four independent experiments. Mean ± SD reported. * p < 0.001

It has been previously reported that the perturbation of KIF1A’s interaction with the C-terminal tail reduces the motor’s landing rate [22]. Measuring KIF1A landing rate on subtilisin-treated microtubules confirms the necessity of this interaction, showing that the perturbation of KIF1A’s interaction with the C-terminal tail reduces its landing rate on the microtubule by 65% in the ADP state (Figure 6, Table 1). Taken together, these results expose KIF1A’s reliance on the microtubule C-terminal tail to engage with the microtubule, initiate pausing behavior, and generate superprocessive overall run lengths.

### Conserved lysine residues of the K-loop are key mediators in KIF1A pausing and motility

Given KIF1A’s reliance on microtubule C-terminal tails to engage in pausing behavior, we next investigated this interaction further by perturbing the electrostatic interaction between the KIF1A K-loop and the microtubule C-terminal tails while maintaining the K-loop structure. First, we assessed the dependence of KIF1A pausing on the local electrostatic environment, by increasing the salt concentration of our motility buffer. In doing so, we observed a decrease in pause duration with no significant changes in pause frequency or any motility parameters (Figure S7). This result lead us to further consider how positively charged residues within the K-loop contribute to KIF1A pausing and motility.

It has been previously reported that mutating all six lysines of the K-loop to alanine residues abolish KIF1A motility and interaction with the microtubule [22]. Therefore, in this complementary approach, we mutated the three most conserved lysines of the kinesin-3 family to maintain a population of KIF1A motors able to engage with the microtubule surface (Figure 4A). KIF1A K-loop mutants (Tri-Ala KIF1A) displayed a marked reduction in both motility and pausing behavior (Figure 4B). Specifically, both the overall run length and processive run length were significantly reduced by 61% and 32%, respectively, when compared to WT KIF1A on taxol-stabilized microtubules (Figure 4C, Table 1). Due to fewer pausing events, there was a 23% increase in overall speed but no change in processive speed (Table 1, Figure S2, S3). Tri-Ala KIF1A mutants also exhibited impaired pausing behavior quantified by a 55% reduction in pause duration and a 75% decrease in pause frequency (Figure 4D, 4E, Table 1). As expected from previous work [22], the Tri-Ala KIF1A mutant also had a dramatic, 92% reduction in landing rate when compared to WT KIF1A on taxol-stabilized microtubules (Figure 6, Table 1). These results further confirm the necessity of the KIF1A K-loop/microtubule C-terminal tail interaction for not only landing rate as previously described [22], but for motility and pausing behavior as well.

**Figure 4.**
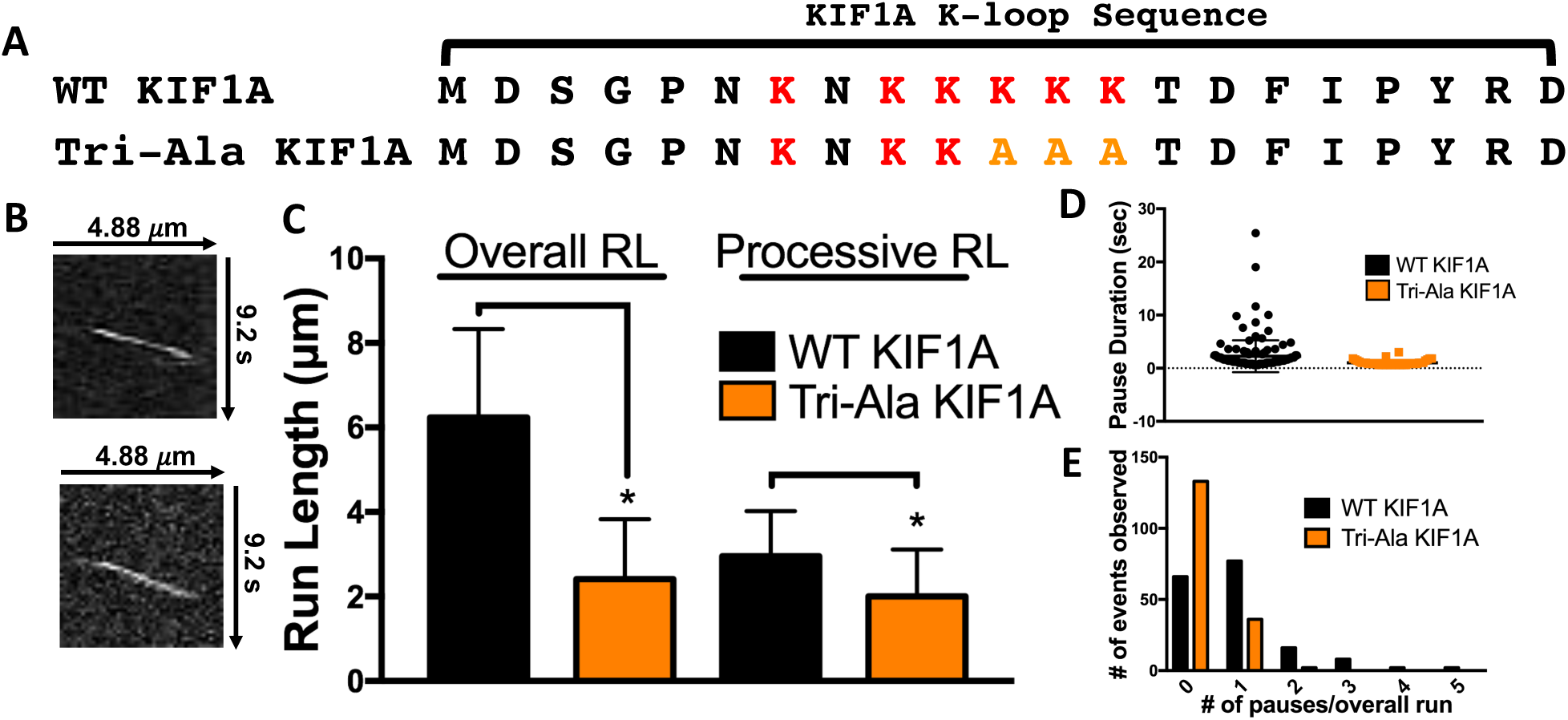
**A)** Amino acid sequence alignment of the KIF1A loop 12 (K-loop) of WT KIF1A and the Tri-Ala KIF1A mutant. For the Tri-Ala mutant, the three most conserved lysines amongst the kinesin-3 family have been mutated to alanine residues (depicted in yellow). **B)** Representative kymographs detailing a significant reduction in pause frequency as well as overall and processive run length of the KIF1A Tri-Ala mutant. **C)** The Tri-Ala mutation of the KIF1A K-loop significantly reduced both overall (6.24 ± 2.09 µm to 2.41 ± 1.42 µm [n=173]) and processive (2.95 ± 1.07 µm to 2.00 ± 1.11 µm [n=207]) run length. **D)** Pause duration decreased from 2.25 ± 2.99 sec (WT KIF1A) to 1.01 ± 0.53 sec (Tri-Ala KIF1A [n=36]) upon mutation of the K-loop. **E)** Mutation of the K-loop significantly decreased pause frequency from 0.95 pauses/overall run (WT KIF1A) to 0.23 pauses/overall run (Tri-Ala KIF1A). Run length values and standard deviations were calculated as previously reported [73]. Kinesin-1 was used as an experimental control across all conditions (Figure S2-S5). Each condition is representative of at least four independent experiments. Mean ± SD reported. * p < 0.001

### Microtubule C-terminal tail polyglutamylation regulates KIF1A pausing behavior and motility

With the KIF1A K-loop/microtubule C-terminal tail interaction characterized, we next aimed to identify a specific component of the microtubule C-terminal tails that could mediate KIF1A pausing and motility. It has been previously established that reduced levels of α-tubulin polyglutamylation results in improper KIF1A localization and insufficient cargo delivery in hippocampal and superior cervical ganglia neurons of ROSA22 mice [21]. As KIF1A is expressed in neurons, we next considered that neuronal microtubules are subject to extensive C-terminal tail polyglutamlation [18], and that this posttranslational modification may be important for KIF1A regulation. To determine the importance of tubulin polyglutamylation on KIF1A behavior and motility, we assessed KIF1A’s pausing and superprocessivity on purified HeLa tubulin known to have markedly less polyglutamylation than neuronal tubulin (Figure 5A) [34-36]. KIF1A overall run length was reduced by 34% on taxol-stabilized HeLa microtubules when compared to taxol-stabilized neuronal microtubules with no significant reduction in processive run length (Figure 5C, Table 1). In regards to pausing behavior, we observed a 58% reduction in pause frequency (Figure 5B, 5E, Table 1) and a 45% decrease in pause duration (Figure 5D, Table 1). Due to the reduction in pausing events, the overall speed of KIF1A on HeLa tubulin increased by 26% when compared to KIF1A on neuronal microtubules, while processive speed decreased by only 7% (Table 1, Figure S2, S3). We also observed a substantial 94% reduction in KIF1A landing rate on HeLa microtubules when compared to neuronal microtubules (Figure 6, Table 1). Taken together, these results confirm that microtubule C-terminal tail polyglutamylation dictates KIF1A’s ability to pause, and mediates superprocessivity.

**Figure 5.**
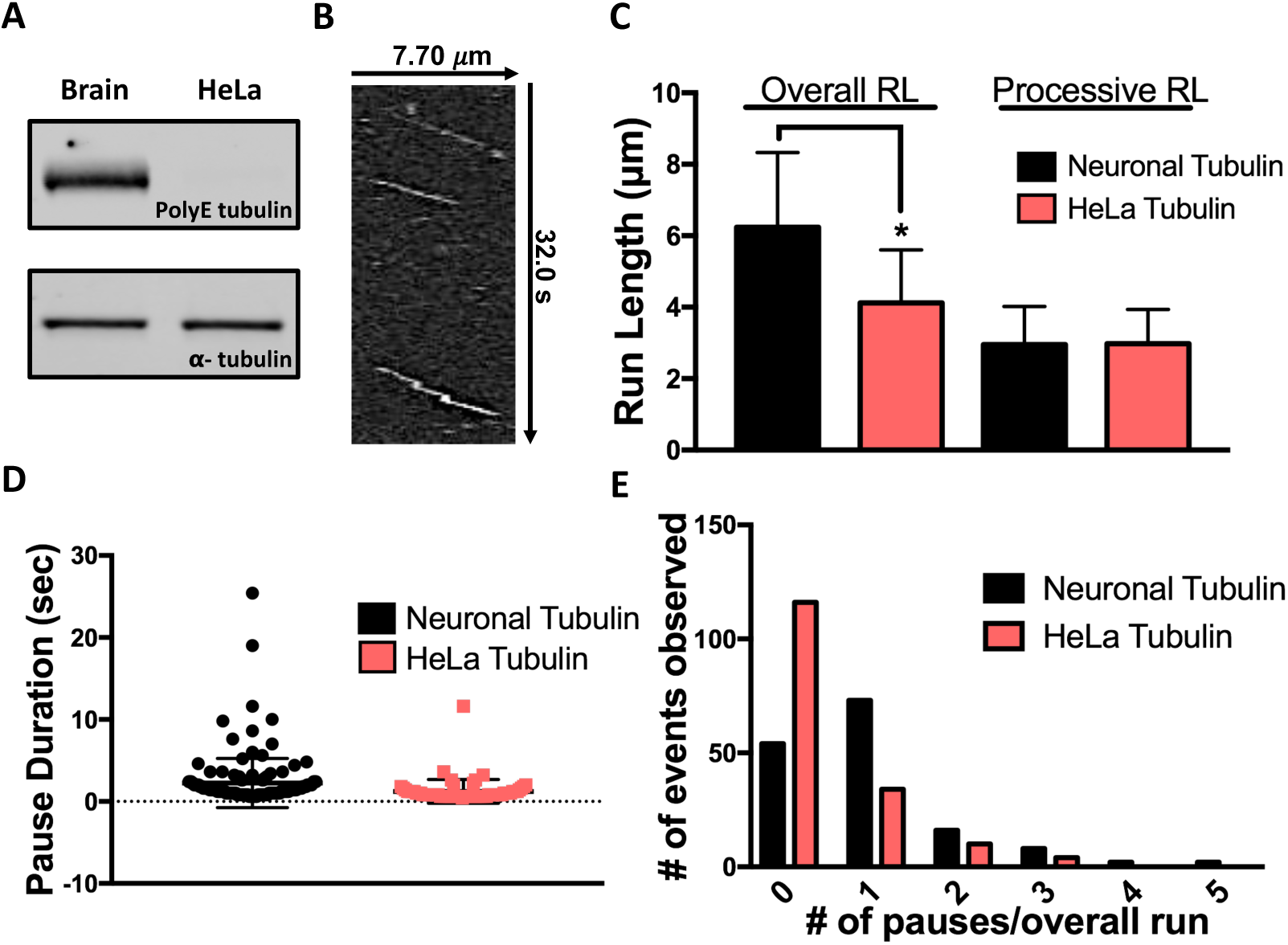
**A)** Western blot revealing higher levels of polygltamylation (polyE) on purified neuronal (brain) tubulin in comparison to purified HeLa tubulin (top) and total α-tubulin loading control (bottom) [34-36]. **B)** Representative kymograph detailing a reduction in pause frequency and run length of KIF1A on HeLa microtubules. **C)** The overall run length of KIF1A was reduced from 6.24 ± 2.09 µm to 4.12 ± 1.48 µm [n=164] on HeLa microtubules, while processive run length was not significantly changed (2.95 ± 1.07 µm and 2.98 ± 0.96 µm [n=222], respectively). **D)** Pause duration decreased from 2.25 ± 2.99 sec to 1.23 ± 1.46 sec [n=58] on HeLa microtubules. **E)** KIF1A pause frequency was significantly decreased from 0.95 pauses/overall run to 0.40 pauses/overall run on HeLa microtubules, when compared to taxol-stabilized neuronal microtubules. Run length values and standard deviations were calculated as previously reported [73]. Kinesin-1 was used as an experimental control across all conditions (Figure S2-S5). Each condition is representative of at least four independent experiments. Mean ± SD reported. * p < 0.001

**Figure 6.**
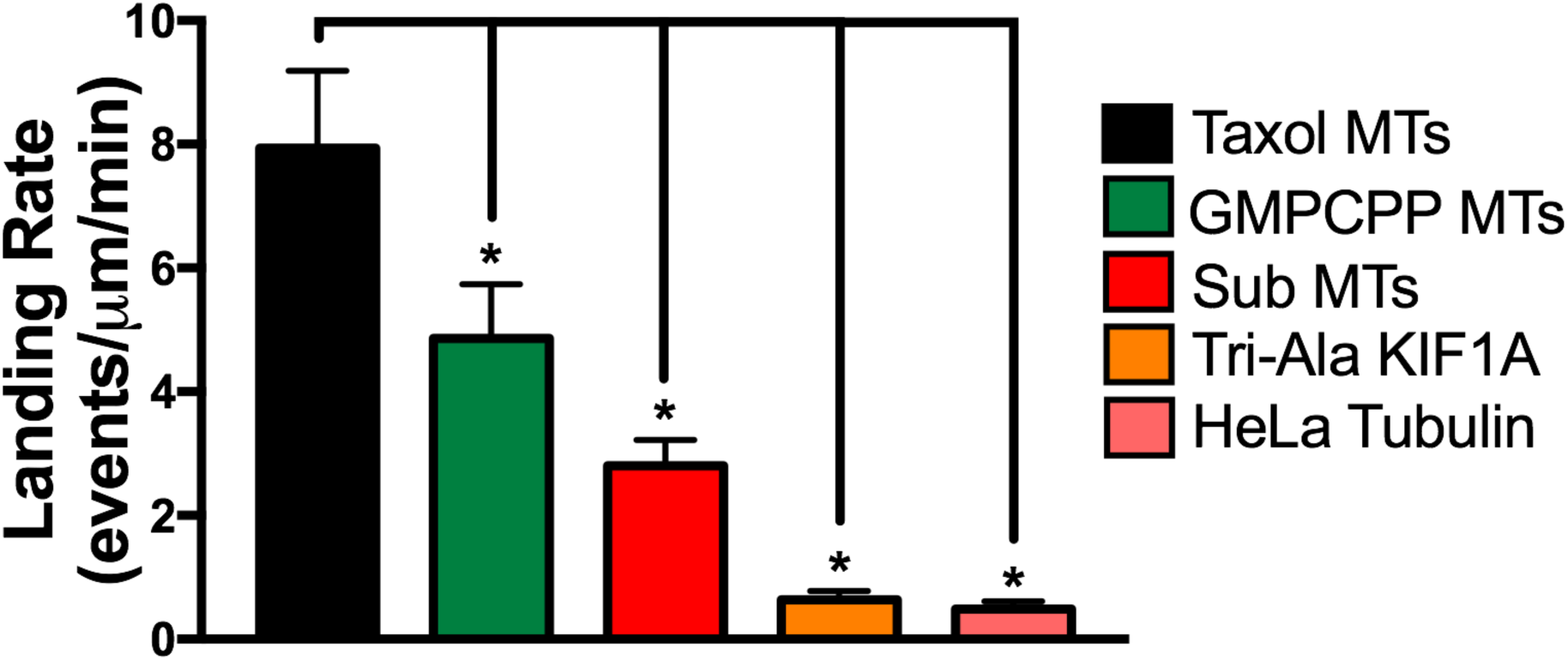
Quantification of KIF1A landing rate in the ADP state on indicated microtubule (MT) subsets (Mean ± SD). On taxol-stabilized microtubules KIF1A had a landing rate of 7.93 ± 1.25 events/μm/min (n=1552). On GMPCPP microtubules, KIF1A landing rate was significantly reduced to 4.86 ± 0.88 events/μm/min (n=958). Removal of microtubule C-terminal tails (Sub MTs) significantly reduced KIF1A landing rate to 2.80 ± 0.42 events/μm/min (n=717). Mutation of the KIF1A K-loop (Tri-Ala KIF1A) significantly reduced the landing rate to 0.64 ± 0.14 events/μm/min (n=527). On HeLa microtubules, KIF1A landing rate was significantly reduced to 0.49 ± 0.13 events/μm/min (n=429). Each condition is representative of at least four independent experiments. Mean ± SD reported. * p < 0.001

## DISCUSSION

The superprocessivity of the kinesin-3 family member KIF1A has been previously established, supporting its known role in long-distance axonal transport [3, 10]. In our study, we present a novel molecular mechanism mediating KIF1A’s superprocessivity. First, we have revealed that KIF1A engages in previously unreported pauses on the microtubule lattice every ∼ 3 μm. Furthermore, we show that these pauses are mediated by interactions with the KIF1A K-loop and polyglutamylated microtubule C-terminal tails. When the K-loop/C-terminal tail interaction is interrupted (i.e., C-terminal tail removal, K-loop mutation, or decreased C-terminal tail polyglutamylation), KIF1A loses its ability to pause on the microtubule lattice. In this scenario, KIF1A can still engage in an initial processive run of ∼ 3 μm but cannot use pauses to string together multiple processive runs to generate a longer overall run. Of note, even in the absence of pauses, KIF1A remains a superprocessive motor, with processive run lengths of ∼ 3 μm, and superprocessivity is enhanced when KIF1A pauses and links processive runs together. KIF1A’s overall speed is also reduced in the absence of pauses, with no effect on processive speed. This response is expected, considering that a reduction in pause frequency will reduce a slower population of motors undergoing pauses. Taken together, our findings confirm our hypothesis that posttranslational microtubule C-terminal tail polyglutamylation is a regulatory mechanism of KIF1A by regulating pausing behavior and subsequent motility.

With a molecular regulatory mechanism identified in regards to KIF1A pausing and motility, a question still remains: *how* is KIF1A able to pause in between processive segments? In this study, we’ve shown that KIF1A walks a superprocessive distance (∼3 μm on average) before initiating a pause, supporting the notion that KIF1A pausing is rare, but observed due to the superprocessive nature of the motor itself. We suspect that pausing is mediated in part by the strength of the E-hook and K-loop electrostatic interaction, supported by conserved motility parameters, as well as decreased pause duration in a higher salt motility buffer (Figure S7). However, to further consider structural interactions between KIF1A and the microtubule during a pause, the catalytic nature of the KIF1A ATP hydrolysis cycle may provide a clue. The KIF1A K-loop (loop 12) and another surface loop of the KIF1A motor domain (loop 11) work together to facilitate KIF1A binding in the transitions between nucleotide states [24]. In support of our data, this process ultimately results with the K-loop interacting with the microtubule C-terminal tail in the ADP state after experiencing a transient binding state in which both loops are extended up away from the microtubule surface, in a confirmation that could facilitate a pause based on C-terminal tail accessibility. This postulation assumes that a pause would happen when both motor domains are in the same nucleotide state at the same time, making this a relatively rare event. To rigorously test how KIF1A pauses is an enticing future direction beyond the scope of this current study.

The connection we present between microtubule C-terminal tail polyglutamylation and KIF1A function is important in advancing our understanding of regulatory mechanism for KIF1A cargo delivery in cells. For example, levels of microtubule C-terminal tail polyglutamylation are highly tuned to the stages of neural development [18, 37]. KIF1A is also a key player in neural development, aiding in autophagy essential for developmental synaptogenesis [38] and facilitating basal interkinetic nuclear migration [39]. Compartmentalizing neuronal tubulin polyglutamylation into localized hotspots [40] may contribute an advantageous level of regulation to fine-tune KIF1A function during these developmental processes. Furthermore, temporally regulating levels of polyglutamylation has been shown to be a potent way to either enhance or reduce the magnitude of KIF1A cargo transport, as supported by the previously mentioned ROSA22 mouse model with impaired polyglutamylation and altered KIF1A cargo trafficking [21]. Taken together with our findings, this mechanism now allows us to propose a molecular link for this physiological impairment, highlighting the need for further studies of how other posttranslational modifications may regulate KIF1A function.

Our discovered mechanism of KIF1A regulation is one way that posttranslationally modifying the microtubule structure regulates the behavior and motility of KIF1A with broader cellular and systemic effects. However, transport of cellular cargo along neuronal microtubules presents many complexities beyond C-terminal tail posttranslational modification, such as obstacles that KIF1A and other kinesin motors must navigate around. The necessity to regulate kinesin cargo transport through pausing is a pre-established concept on a cellular level, as many neuronal cargo have been observed to halt processive movement in order to navigate around obstacles in the crowded cellular environment [41-44]. One group of obstacles KIF1A will encounter are microtubule associated proteins (MAPs) that bind to the microtubule surface and regulate microtubule motor based cargo transport. Microtubule affinity of MAPs such as MAP1A, MAP1B, MAP2, and Tau are all known to be regulated by levels of microtubule C-terminal tail polyglutamylation [45, 46]. Tau, an axonal-specific MAP known to regulate the motility of kinesin-1 family members [47-49], interacts with tubulin C-terminal tails in a unique diffusive behavior that is important for Tau function on a cellular and systemic level [50]. With Tau competing with KIF1A for microtubule C-terminal tails, our model would suggest that Tau is capable of regulating KIF1A pausing and motility. This concept, supported by recent compelling work on the influence of MAPs and septins on KIF1A function [32, 51-53], is a compelling topic for further study.

The importance of KIF1A motility regulation is critical when one considers the extensive range of axonal cargo that KIF1A transports along neuronal microtubules [3, 9, 11, 54-60]. Furthermore, until we advance our understanding of KIF1A regulation, we are limited in our ability to address the disease state manifestations of altered KIF1A function. The regulatory capabilities of Tau’s binding behavior on KIF1A cargo transport are critical to understanding KIF1A’s role in in diseases known for impaired axonal transport, such as frontotemporal dementia (FTD), where changes in Tau isoform expression have been correlated to disease progression [61]. For example, pathological overexpression [62-64] of the more diffusive Tau isoforms [65] and reduced transport of cargo known to be transported by KIF1A are observed in FTD [11, 12, 66]. As we speculate that the diffusive binding state of Tau may compete with KIF1A for the microtubule C-terminal tail, this is an enticing concept for further investigation on a molecular and cellular levels. In summary, our discovery of a microtubule C-terminal tail polyglutamylation-mediated mechanism for regulation of KIF1A motility provides crucial insight as to how KIF1A cargo delivery is regulated during axonal transport and how this process may become altered in the disease state.

## METHODS

### Tubulin isolation, microtubule preparation and labelling

Neuronal tubulin was isolated from bovine brains donated from Vermont Livestock Slaughter & Processing (Ferrisburgh, VT) using a high molarity PIPES buffer (1 M PIPES, pH 6.9 at room temperature, 10 mM MgCl_2_, and 20 mM EGTA) as previously described [67]. Purified tubulin was clarified using ultracentrifugation for 20 minutes at 95,000 rpm at 4° C in an Optima TLX Ultracentrifuge (Beckman, Pasadena, CA). After clarification, tubulin concentration was calculated using the tubulin extinction coefficient of 115,000 cm^-1^ M^-1^ and read at 280nm in the spectrophotometer.

Tubulin was purified from HeLa Kyoto cells with TOG affinity column chromatography using gravity-flow setup [68, 69]. Cells were resuspended in BRB80 (80 mM PIPES, 1 mM EGTA, 1 mM MgCl_2_ pH 6.8) supplemented with 1 mM DTT and protease inhibitors (cOmplete*™* Mini, EDTA-free [Sigma-Aldrich, St. Louis, MO] and 0.2 mM PMSF) and sonicated. Cleared lysate was loaded onto a TOG column, in which approximately 15 mg of bacterially purified GST-TOG1/2 protein was conjugated with 1 ml of NHS-activated Sepharose 4 Fast Flow resin (GE Healthcare; Marlborough, MA). Wash, elution, desalting and concentration was carried out as described in Hotta et al, 2016 [69]. Glycerol was not added to the purified tubulin. Aliquoted tubulin was snap frozen in liquid nitrogen and stored at -80°C.

Clarified tubulin was supplemented with 1mM GTP (Sigma-Aldrich) or guanosine-5′-[(*α*,*β*)-methyleno]triphosphate sodium salt (GMPCPP; Jena Bioscience, Jena, Germany). To label microtubules, unlabeled tubulin was mixed with rhodamine-labeled tubulin (Cytoskeleton, Denver, CO) at a ratio of 100:1. Microtubules were stabilized with GMPCPP following previously reported methods[70] or polymerized at 37° C for 20 minutes and stabilized with 20 μM paclitaxel (taxol; Sigma-Aldrich) in DMSO. Microtubules were diluted to a working concentration of 1 μM in P12 Buffer (12mM PIPES, 1mM MgCl_2_, 1 mM EGTA, pH 6.8 supplemented with 20 μM paclitaxel) [22].

### Plasmids, mutagenesis, and cell lysate motor expression

Mammalian KIF1A(1-393)-LZ-3xmCitrine plasmid was a generous gift of Kristen Verhey (University of Michigan, Ann Arbor, MI). KIF5C(1-559)-GFP and KIF1A(1-396)-LZ-GFP mammalian plasmids were a gift from Gary Banker (Oregon Health & Science University). TriAla-KIF1A-LZ-3xmCitrine was generated using the QuikChange II XL Site-Directed Mutagenesis Kit (Agilent Technologies, Santa Clara, CA). COS-7 monkey kidney fibroblasts (American Type Culture Collection, Mansses, VA) were cultured in DMEM-GlutaMAX^*™*^ with 10% fetal bovine serum (FBS) at 37° C with 5% CO_2_. Cells were transfected with 1μg of either WT KIF1A-LZ-3xmCitrine, Tri-Ala KIF1A-LZ-3xmCitrine, or KIF1A-GFP mammalian plasmid using the Lipofectamine 2000 delivery system (Thermo Fisher Scientific, Waltham, MA) and incubated in Opti-MEM^TM^ (Thermo Fisher Scientific) media with 4% FBS. The next day, cells were harvested, pelleted, and washed with DMEM GlutaMAX^*x*^. The pellet was vigorously resuspended in lysis buffer (25mM HEPES, 11mM K^+^ Acetate, 5mM Na^+^ Acetate, 5mM MgCl_2_, 0.5mM EGTA, 1% Triton X-100, 1mM PMSF, 1mg/ml pepstatin, 10 μg/ml leupeptin, 5μg/ml aprotinin) and centrifuged at room temperature for 15 minutes at 14,000 rpm. Relative amounts of protein between preps and motor constructs was determined using densitometry, and supernatant containing expressed motor protein was aliquoted and stored at -80° C until further use.

### Subtilisin treatment

To remove C-terminal tails, 5μM paxlitaxel stabilized microtubules were treated with 0.05μM Subtilisin A (Sigma Aldrich), resuspended in P12 Buffer, for 45 minutes at 25° C. The reaction was stopped by the addition of 5mM phenylmethanesulfonyl fluoride. Subtilisin treatment was confirmed by Coomassie staining of SDS-PAGE denaturing gel. Tubulin concentration was determined as previously described.

### In vitro single-molecule TIRF

Flow chambers used in *in vitro* TIRF experiments were constructed as previously described [49]. Flow chambers were incubated with monoclonal anti-β III (neuronal) antibodies at 33 μg/ml for 5 minutes, then washed twice with 0.5mg/ml bovine serum albumin (BSA; Sigma Aldrich) and incubated for 2 minutes. 1 μM of microtubules (any experimental condition) were administered and incubated for 8 minutes. Non-adherent microtubules were removed with a P12 wash supplemented with 20μM paclitaxel. Kinesin motors in Motility Buffer consistent with past literature (MB; 12mM PIPES, 1mM MgCl_2_, 1 mM EGTA, supplemented with 20 μM paclitaxel, 10 mM DTT, 10 mg/ml BSA, 2 mM ATP and an oxygen scavenger system [5.8 mg/ml glucose, 0.045 mg/ml catalase, and 0.067 mg/ml glucose oxidase; Sigma Aldrich]) [22], or a High Salt Motility Buffer (same as Motility Buffer, with the only change being PIPES concentration raised to 80mM), supplemented with 2mM ATP, were added to the flow cell just before image acquisition. For landing rate assays, *in vitro* KIF1A motility assays were prepared as described above, with the only change being that MB was supplemented with 2mM ADP. Control *Drosophila melanogaster* biotin-tagged kinesin-1 motors were labeled with streptavidin-conjugated Qdot 655 (Life Technologies, Carlsbad, CA) at a 1:4 motor:Qdot ratio as previously described [49, 71].

Total internal reflection fluorescence (TIRF) microscopy was performed at room temperature using an inverted Eclipse Ti-E microscope (Nikon, Melville, NY) with a 100x Apo TIRF objective lens (1.49 N.A.) and dual iXon Ultra Electron Multiplying CCD cameras, running NIS Elements version 4.51.01. Rhodamine-labeled microtubules were excited with a 561 and a 590/50 filter. KIF1A-LZ-3xmCitrine (WT or Tri-Ala) motors were excited with a 488 laser and a 525/50 filter. Qdot 655 conjugated kinesin-1 motors were excited with a 640 laser and 655 filter. All movies were recorded with an acquisition time of 200 msec for 500 frames (100 sec observation).

### Photobleaching assay

As an additional control, a photobleaching assay was performed to confirm that KIF1A run lengths were not underestimated due to photobleaching of the C-terminal 3xmCitrine tag (Figure S2). Motors were adhered to microtubules in the presense of adenylylimidodiphosphate (AMPPNP; Sigma-Aldrich) and washed once to remove unattached motors. Using Image J, a region of interest (ROI) was drawn around each fluorescent spot and the average intensity of each pixel within the ROI was measured over time. Intensity was background corrected and a motor was considered photobleached when the ROI average intensity was 0.

### Western blot analysis

Purified tubulin protein was separated by electrophoresis on Mini-PROTEAN^*®*^ TGX*™*gels (Bio-Rad, Hercules, CA) and transferred to a polyvinylidene fluoride membrane (Bio-Rad). Membranes were blocked in in a 1:1 solution of phosphate-buffered saline (PBS; 155 mM NaCl, 3 mM Na_2_HPO_4_, and 1 mM KH_2_PO_4_, pH 7.4) and Odyssey^*®*^ blocking reagent (Li-COR, Lincoln, NE). A mouse anti-polyglutamylated tubulin antibody (1:8,000; GT335, AdipoGen, San Diego, CA) or a mouse anti-alpha tubulin antibody (1:10,000; DM1A, Sigma-Aldrich) was administered to membrane, followed by a secondary DyLight 800 anti-mouse IgG antibody (1:10,000; Thermo Fisher Scientific). Secondary antibody fluorescence was detected using an Odyssey CLx (Li-COR).

### Data analysis

Motility events were analyzed as previously reported [71, 72]). In brief, overall run length motility data was measured using the ImageJ MTrackJ plug-in, for a frame-by-frame quantification of KIF1A motility. Pauses are defined as segments in where the average velocity is less than 0.2 μm/s over three frames or more. Processive events were identified at the boundaries of pausing events. Average overall/processive speed were plotted as a histogram, with mean and SD are reported in Table 1.

Kymographs of motor motility were created using the MultipleKymograph ImageJ plug-in, with a set line thickness of 3. To correct for microtubule track length effects on motor motility, overall/processive run length data was resampled to generate cumulative frequency plots (99% CI), using previously reported methods [73]. After data resampling, statistical significance between motor run length and speed data sets was determined using a paired t-test.

To determine the landing rate of KIF1A on various microtubule conditions, the total number of motor landing events on the microtubule was divided by the length of the microtubule and further divided by the duration of the movie (events/μm/minute). To be considered a landing rate event, a motor must remain on the microtubule for three consecutive frames under ADP state conditions.

## ACKNOWLEDGEMENTS

We thank David Warshaw and Guy Kennedy for training and use of the TIRF microscope at the University of Vermont. A special thank you to Vermont Livestock Slaughter & Processing (Ferrisburgh, VT) for supporting our work. We thank Gary Banker & Marvin Bentley for the gifted pBa.Kif1a 1-396.GFP was a gift from (Addgene plasmid # 45058), William Hancock for the use of his kinesin-1 construct, and Adam Hendricks for support. This work was supported by National Institute of General Medicine Sciences/National Institutes of Health funding to C.B. (Grant GM101066) and R.O. (GM086610).

## AUTHOR CONTRIBUTIONS

Experiments were conceived and designed by DVL, CLB, and KJV. Experiments were performed and analyzed by DVL, OJZ and TH. Manuscript was drafted by DVL and CLB. TH and RO contributed reagents. Manuscript was revised and given final approval by all authors.

